# Tobacco exposure associated with oral microbiota oxygen utilization in the New York City Health and Nutrition Examination Study

**DOI:** 10.1101/470286

**Authors:** Francesco Beghini, Audrey Renson, Christine P. Zolnik, Ludwig Geistlinger, Mykhaylo Usyk, Thomas U. Moody, Lorna Thorpe, Jennifer B. Dowd, Robert Burk, Nicola Segata, Heidi E. Jones, Levi Waldron

## Abstract

**Purpose:** The effect of tobacco exposure on the oral microbiome has not been established.

**Methods:** We performed amplicon sequencing of the 16S ribosomal RNA gene V4 variable region to estimate bacterial community characteristics in 259 oral rinse samples, selected based on self-reported smoking and serum cotinine levels, from the 2013-14 New York City Health and Nutrition Examination Study. We identified differentially abundant operational taxonomic units (OTUs) by primary and secondhand tobacco exposure, and employed “microbe set enrichment analysis” to assess shifts in microbial oxygen utilization.

**Results:** Cigarette smoking was associated with depletion of aerobic OTUs (Enrichment Score test statistic ES = −0.75, p = 0.002) with a minority (29%) of aerobic OTUs enriched in current smokers compared to never smokers. Consistent shifts in the microbiota were observed for current cigarette smokers as for non-smokers with secondhand exposure as measured by serum cotinine levels. Differential abundance findings were similar in crude and adjusted analyses.

**Conclusion:** Results support a plausible link between tobacco exposure and shifts in the oral microbiome at the population level through three lines of evidence: 1) a shift in microbiota oxygen utilization associated with primary tobacco smoke exposure, 2) consistency of abundance fold-changes associated with current smoking and shifts along the gradient of secondhand smoke exposure among non-smokers, and 3) consistency after adjusting for *a priori* hypothesized confounders.

**Highlights:**

- Cigarette smoke was associated with microbial anaerobiosis in oral rinse specimens
- The microbiome shifts associated with smoking and secondhand exposure were correlated
- Shifts in oral bacterial oxygen utilization may mediate smoking and health outcomes
- We propose “microbe set enrichment analysis” for interpreting shifts in the microbiome

## Introduction

Dysbiosis of the dental plaque microbiome is a necessary step in the etiology of periodontitis and caries [1], which have been linked to systemic illness, including cardiovascular diseases [2], type 2 diabetes mellitus [3], obesity [4], low birth weight and preterm birth [5], rheumatoid arthritis [6], chronic obstructive pulmonary disease [7], and oral and digestive cancers [8]. Tobacco exposure is a cause of these outcomes [9–14], but whether it causes them through shifts in the general oral microbiome is unknown [15]. If tobacco smoke causes harmful alterations of the oral microbiome, interventions targeting the oral microbiome could mitigate the impact of tobacco exposures. A key aspect of making this distinction lies in establishing whether a range of tobacco exposures, including cigarette smoking, secondhand smoke exposure, hookah and e-cigarette use, cause substantial changes in the structure and function of the general oral microbiome.

Evidence suggests that tobacco smoke exposure causes alterations to the oral microbiome, selecting a community enriched with opportunistic pathogens [16,17] and negatively impacting the resilience and colonization resistance of the sub- and supragingival biofilms [18]. Such alterations may occur directly due to selective toxicity [19], or indirectly via alteration of the host immune system to produce both pro- and antiinflammatory effects [20–22] which alter the oral biofilm and mucosal microbial habitats. Another potential mechanism by which tobacco smoke reconfigures the oral microbiome is via depletion of oxygen [23], creating a hypoxic oral environment that favors anaerobiosis. Tobacco smoke may also favor anaerobiosis by increasing the amount of free iron [24], and inhibiting oral peroxidase [25]. Anaerobic glycolysis in human salivary cells has been shown to dramatically increase after exposure to tobacco smoke [26], and human experiments show reduction in periodontal pocket oxygen tension [27] and redox potential [23] after smoking cigarettes. Low throughput studies of the oral microbiome have shown greater abundance of the anaerobes *Prevotella intermedia* [28] and *Lactobacillus spp*. [29] in cigarette smokers. Ad hoc findings from high throughput studies have suggested that smokers have greater abundance of anaerobic microorganisms [30] and depletion of microbial functional pathways related to aerobic respiration [15]. Thus tobacco exposure could plausibly cause changes to the oral microbiome, but available results are limited to laboratory and small-sample studies.

This study suggests a causal link between tobacco exposure and alterations to the saliva microbiome among participants of the 2013-14 New York City Health and Nutrition Examination Study (NYC HANES). It contrasts current smokers to non-smokers with no recent secondhand exposure, and investigates a dose-response relationship among non-smokers with varying degrees of secondhand exposure assessed by quantitative serum cotinine level. It further investigates former smokers and smokers of e-cigarettes and hookah. Multiple lines of causal inference are used to test the hypothesis that tobacco smoke alters the saliva microbiome: controlling for hypothesized confounders, testing for a dose-response relationship, and testing for altered oxygen requirements of the microbial communities associated with tobacco exposure.

## Materials and Methods

### 2013-2014 NYC-HANES

Data for the current study are sub-sampled from the 2013-14 NYC HANES, a population-based study of 1,575 non-institutionalized adults in New York City [31] modeled after the United States Centers for Disease Control and Prevention’s National Health and Nutrition Examination Survey (NHANES) [32]. The NYC HANES sample was recruited using a three stage cluster household probabilistic design of all non-institutionalized adults 18 years of age or older. Consenting individuals provided information on smoking, including use of alternative tobacco products such as e-cigarettes and hookahs in the last 5 days, socio-demographic characteristics, and oral hygiene practices through face-to-face interviewing and audio computer assisted interviewing for sensitive questions. Participants also underwent physical examination and provided blood and oral rinse specimens for biomarker analysis. Serum specimens were analyzed for cotinine by liquid chromatography/tandem mass spectrometry [33].

The study was conducted by the City University of New York (CUNY) School of Public Health in collaboration with the New York City Department of Health and Mental Hygiene (NYC DOHMH), with ethical approval from their respective institutional review boards (IRBs). The current sub-study received separate IRB approval from the CUNY School of Public Health.

### Tobacco exposure outcome measures and selection of sub-sample

We selected a sub-sample of 297 participants for oral microbiome assessment based on self-reported tobacco use and serum blood cotinine level, classified into five mutually exclusive groups:

- *Current smokers* (n=90) were selected from participants who reported smoking more than 100 cigarettes in their lifetime, smoking a cigarette in the last 5 days, and not using any alternative tobacco product in the last 5 days (the 90 with highest measured serum cotinine were selected).
- *Never smokers* (n=45) were randomly selected from those reporting lifetime smoking of less than 100 cigarettes, no usage of any tobacco product in the last 5 days, and serum cotinine level less than 0.05 ng/mL.
- *Former smokers* (n=45) were randomly selected from participants reporting lifetime smoking of more than 100 cigarettes, but currently not smoking, no use of any tobacco product in the last 5 days, and serum cotinine level less than 0.05 ng/mL.
- *Non-smokers with secondhand exposure* (n=38) comprised all available former or never smokers with serum cotinine between 1 and 14 ng/mL [34]. This group includes four individuals with serum cotinine > 10 ng/mL [35], who may have been light smokers; sensitivity analyses excluding these four had negligible effect (not shown, included in Reproducible Analysis code), so they are included in the secondhand exposure group.
- *“Alternative” smokers* (n=79) were participants with self-reported usage of hookah, cigar, cigarillo and/or e-cigarette in the last 5 days.

For quality control, 5% of samples were randomly selected and sequenced as technical replicates. Results from replicates were used instead of the original sample if the sequencing read count was greater than the original. An additional eight samples failed PCR amplification and were repeated. Fifteen specimens (n=4 current cigarette smokers, n=2 never smokers, n=2 former smokers, n=7 alternative smokers) were discarded for sequencing quality control (below). We excluded an additional 23 participants from the alternative smoker group who also reported smoking cigarettes in the last 5 days, in an attempt to isolate the effect of alternative tobacco exposures, resulting in an analytic sample of 259 participants.

### Specimen collection, processing, and sequence analysis

Specimen collection, processing, and sequence analysis methods are described in detail in a companion paper [36]. In brief, participants were asked to fast for 9 hours prior to oral rinse and blood specimen collection. A 20-second oral rinse was divided into two 5-second swish and 5-second gargle sessions using 15 mLs of Scope^®^ mouthwash, which was transported on dry ice and stored at −80°C. A modified protocol of the QIAamp DNA mini kit (QIAGEN) was used for DNA extraction [36]. DNA was amplified for the V4 variable region of the 16S rRNA gene [37,38]. High-throughput amplicon sequencing was conducted on a MiSeq (Illumina, San Diego, CA) using 2×300 paired-end fragments. 16S read analysis was carried out using QIIME version 1.9.1 [39] and Phyloseq [40]. Paired-end reads were merged with fastq-join [41] and resulting low quality reads (PHRED score < 30) were discarded when joining the split reads (qiime split_libraries_fastq.py). Operational Taxonomic Unit (OTU) picking was performed with an open reference approach by clustering using UCLUST at 97% sequence similarity and taxonomy was assigned using the SILVA 123 [42] database as reference. The QIIME generated OTU table was converted for phyloseq processing. Samples with fewer than 1000 reads (n=15) were removed from the OTU table in the phyloseq preprocessing step. Genera present with a mean relative abundance of less than 2 × 10^−4^ were collapsed as “Other.”

### Unsupervised clustering

We explored differences in beta diversity measures via Principal Coordinates Analysis (PCoA) plots on Weighted UniFrac distances. The grouping of distances by smoking status was tested by PERMANOVA as implemented in the ‘vegan’ package [43], with 999 permutations.

### Oral microbiome measures

We compared oral bacterial community characteristics by tobacco exposure group using four types of oral microbiome measures: 1. alpha (within-sample) diversity of the OTUs present; 2. beta (between-sample) diversity of OTUs; 3. OTU counts at the genus level; 4. enrichment of differential abundance categorized by oxygen requirement. For diversity measures, we estimated Chao1 Index, Shannon Index and observed OTUs, and weighted UniFrac [44] beta diversity using the estimate_richness and distance methods of the phyloseq Bioconductor package [40].

### Differential abundance analysis

We performed crude and adjusted negative binomial log-linear regression of tobacco exposure group to identify differentially abundant OTUs using edgeR [45] Bioconductor package. Low-prevalence OTUs, those without 3 or more reads observed in at least 8 samples, were discarded. For adjusted models, *a priori* hypothesized confounders included age, sex, race, self-reported physical activity, education, diabetes status (based on serum HbA1c), and self-reported gum disease. Education [46,47], age, and sex [48–50] are known to be associated with smoking and could plausibly be associated with oral microbiome characteristics. Race was also treated as a possible confounder as studies suggest differences in nicotine metabolism by race/ethnicity [51]. A False Discovery Rate [FDR, 52] less than 0.05 was considered statistically significant. Results from edgeR were compared to results obtained from the application of DESeq2 [53]. Crude and adjusted coefficients were compared to assess which hypothesized confounders had the greatest effect on adjusted analyses.

### Microbe set enrichment analysis for oxygen requirements

We categorized genera as aerobic, anaerobic, or facultative anaerobic [54] integrating information from the IMG/MER database [55] and from the Bergey’s Manual of Systematics of Archaea and Bacteria [54]. This resulted in three “microbe sets” of OTUs with common oxygen requirements. We applied two concurrent approaches to analyze whether the three microbe sets show coherent changes in abundance of the contained microbes for (i) smokers vs. never smokers (with no recent secondhand smoke exposure), and (ii) among non-smokers with exposure to secondhand smoke. First, over-representation of differentially abundant OTUs in each microbe set was tested based on the hypergeometric distribution (corresponds to a one-sided Fisher’s exact test). Second, Gene Set Enrichment Analysis [GSEA, 56] was used to test whether microbes of a particular microbe set accumulate at the top or bottom of the full OTU vector ordered by direction and magnitude of abundance change between the tested sample groups. Over-representation analysis (ORA) and GSEA were applied as implemented in the EnrichmentBrowser R/Bioconductor package [57]. Application of GSEA incorporated the voom-transformation [58] of OTU counts to concur with GSEA’s assumption of roughly normally distributed data. As the implementations of GSEA and ORA required a binary outcome, serum cotinine levels were binned to contrast the upper tertile (> 4.42 ng/ml) against the lower tertile (< 1.76 ng/ml). We also analyzed serum cotinine level as a continuous measure using Gene Set Variation Analysis [GSVA, 59].

### Statement of reproducible research

Analyses were performed in QIIME version 1.9.1 and R version 3.5.1. All results presented in this manuscript are reproducible by installing the package and compiling its associated vignettes provided at https://github.com/waldronlab/nychanesmicrobiome.

## Results

A total of 1.4 M reads (mean±sd: 4,758±3,463 reads/sample) were generated from 297 saliva mouthwash specimens [31] of NYC-HANES participants selected based on questionnaire and serum cotinine levels (Table 1, with serum cotinine levels by exposure group shown in Supplementary Figure 1). After quality control and filtering, we retained 91.7% of reads (5,007 mean, 3,491 s.d), which were then classified using the QIIME pipeline [39] into 1291 OTUs with more than 10 reads.

**Table 1.**
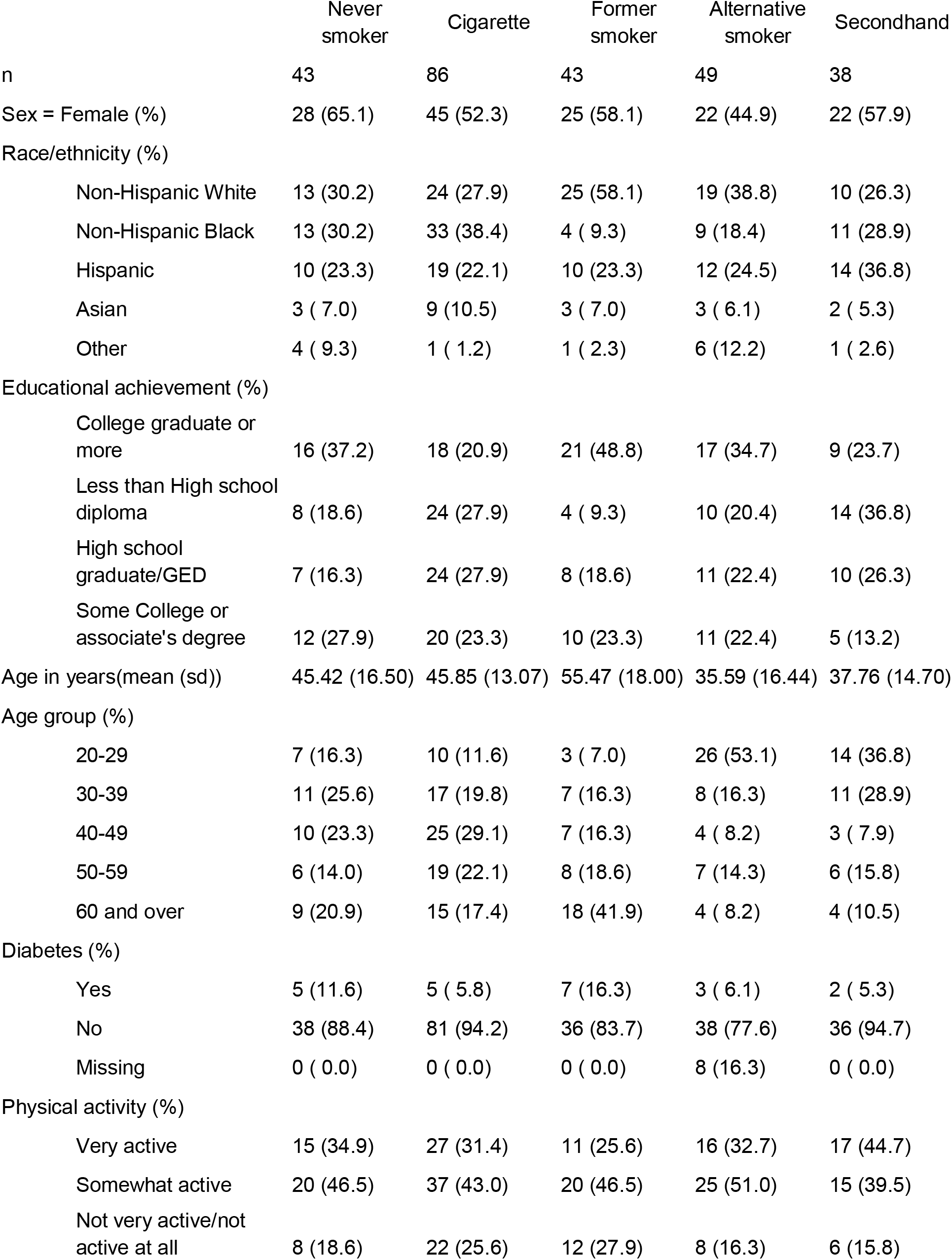
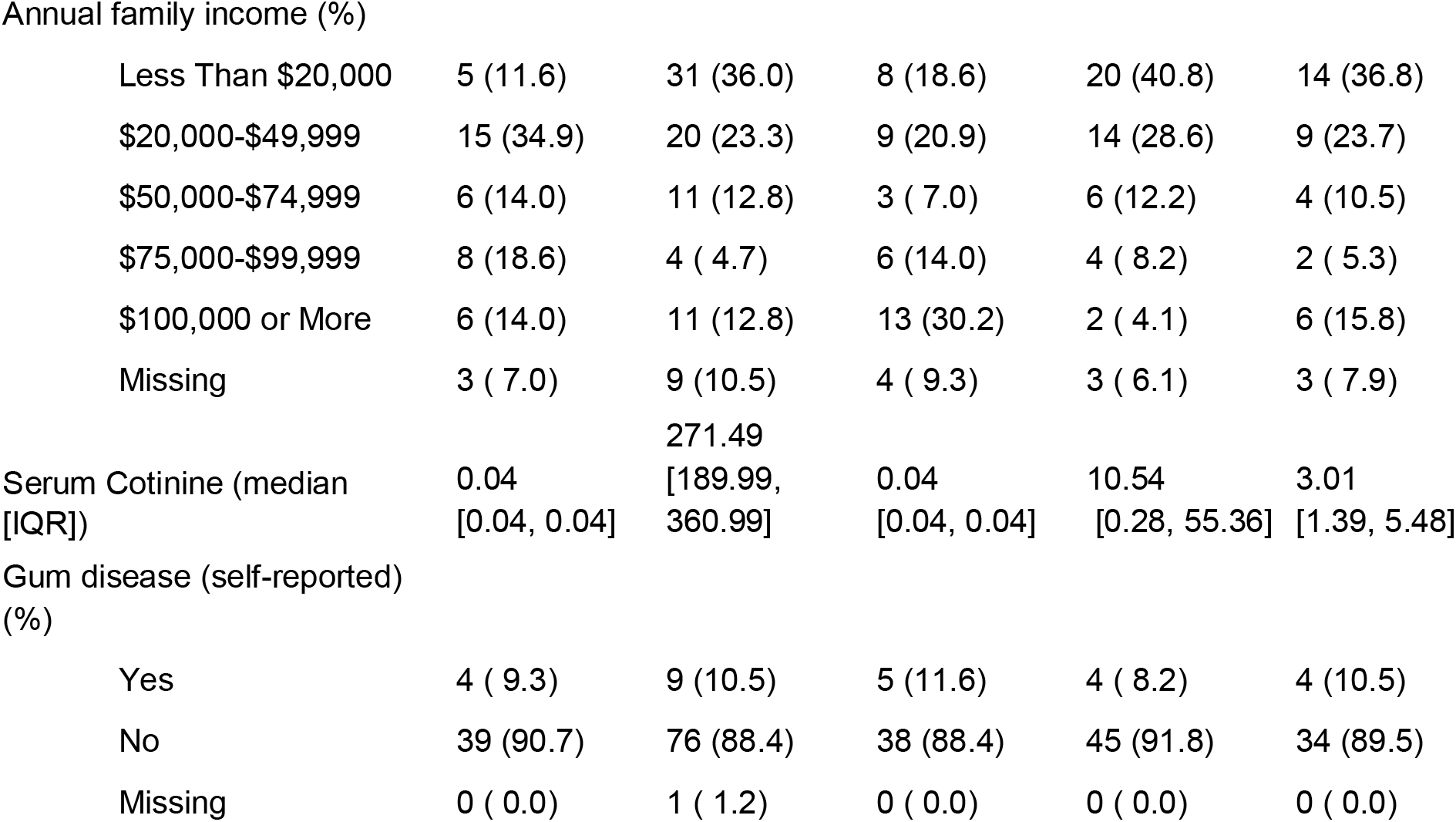
Demographics and characteristics of participants in the 2013-2014 NYC-HANES smoking and oral microbiome study

Taxonomic composition of the final analytic sample (n=259) was predominated by *Streptococcus* (36% average relative abundance) and *Prevotella* (17% average relative abundance), which were present in every sample. Other genera commonly associated with the oral cavity like *Rothia, Neisseria, Veillonella* and *Gemella* were also found with average relative abundances less than 10%.

### Alpha and beta diversity of the oral microbiome by tobacco exposure

Alpha diversities were not significantly different between the five tobacco exposure groups (Shannon Index p=0.95, Observed OTUs p=0.08, Chao1 Index p=0.26 ANOVA test, Supplementary Figure 2). However, beta diversity differed between current cigarette smokers and never smokers, and was larger than differences by race/ethnicity, age, and other sociodemographic measures (Table 2). The overall microbiome composition and structure in these two classes differed and beta diversity was significantly explained by smoking status (R^2^=0.051, p<0.001, PERMANOVA test, Figure 1). Additionally, former smokers were significantly different from current smokers (p=0.001, R^2^=0.044, PERMANOVA test), but not from never smokers (p=0.16, R^2^=0.018, PERMANOVA test). Within former smokers, we found no evidence of differences between those who quit recently versus longer ago (Supplementary Figure 3).

**Figure 1:**
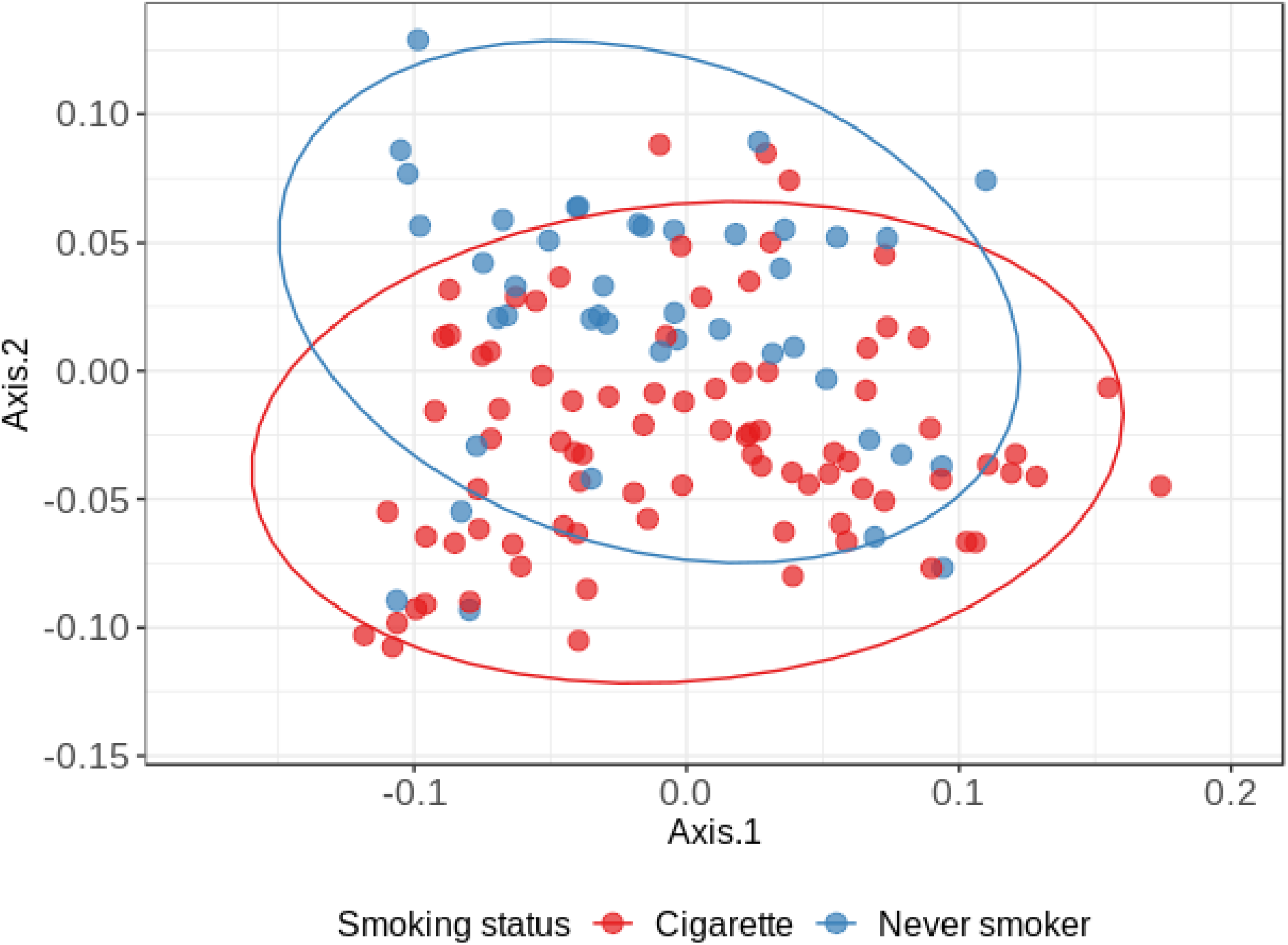
Principal coordinates analysis based on the weighted UniFrac distance. Dots in the ordination plot are samples from never smokers with negligible serum cotinine (blue, n=43) and current cigarette smokers (red, n=86); ellipsis indicating where 95% of observations are expected for each group. A separation between cigarette smokers and never smokers is present and is statistically significant (R^2^ = 0.051, PERMANOVA p<0.001). A gradient also exists for the entire sample (n=259) by measured continuous serum cotinine level (R^2^ =0.0485, PERMANOVA p = 0.001).

**Table 2:** PERMANOVA analysis on Weighted UniFrac distance measure. Model included smoking status (cigarette smokers/never smokers) and other sociodemographic measures. Df: degrees of freedom, R^2^: Coefficient of Determination

|  | Df | F.Model | R<sup>2</sup> | Pr(>F) |
| --- | --- | --- | --- | --- |
| Smoking status<br>(Cigarette smokers vs Never Smokers) | 1 | 7.0845 | 0.05137 | 0.002 |
| Self reported gum disease | 2 | 0.8649 | 0.01254 | 0.496 |
| Race/ethnicity | 4 | 1.6281 | 0.04723 | 0.05 |
| Sex | 1 | 2.2591 | 0.01638 | 0.06 |
| Age groups | 1 | 2.9418 | 0.02133 | 0.014 |
| Physical activity | 2 | 0.8085 | 0.01173 | 0.623 |
| Education level | 2 | 0.7274 | 0.01055 | 0.708 |
| Diabetes | 1 | 0.304 | 0.0022 | 0.937 |

### *Proteobacteria* less abundant in the microbiome of smokers

In crude analyses, 46 differentially abundant OTUs, taxonomically assigned to 28 different genera, were identified between current cigarette smokers and never smokers (Supplementary Figure 4). Relative abundance of OTUs annotated as phylum Proteobacteria *(Neisseria, Lautropia, Haemophilus* and *Actinobacillus)* and Candidate division SR1 were found to be lower in current cigarette smokers compared to never smokers (Proteobacteria phylum t-test p-value=5e-07, logFC=-0.84, Supplementary Figure 5).

Adjusted differential abundance analysis, accounting for hypothesized confounders, identified fewer (n=21) differentially abundant OTUs between current and never smokers (Figure 2). The phylum Proteobacteria was still identified as less-abundant in current smokers in the adjusted model (t-test p-value=8e-07, logFC=-0.85). Adjusted coefficients were slightly attenuated toward the null compared to crude estimates (Figure 3). Addition of one hypothesized confounder at a time showed that age and education had the strongest impact, resulting in a median decrease in coefficient magnitude of 2 and 3 percent, respectively.

**Figure 2:**
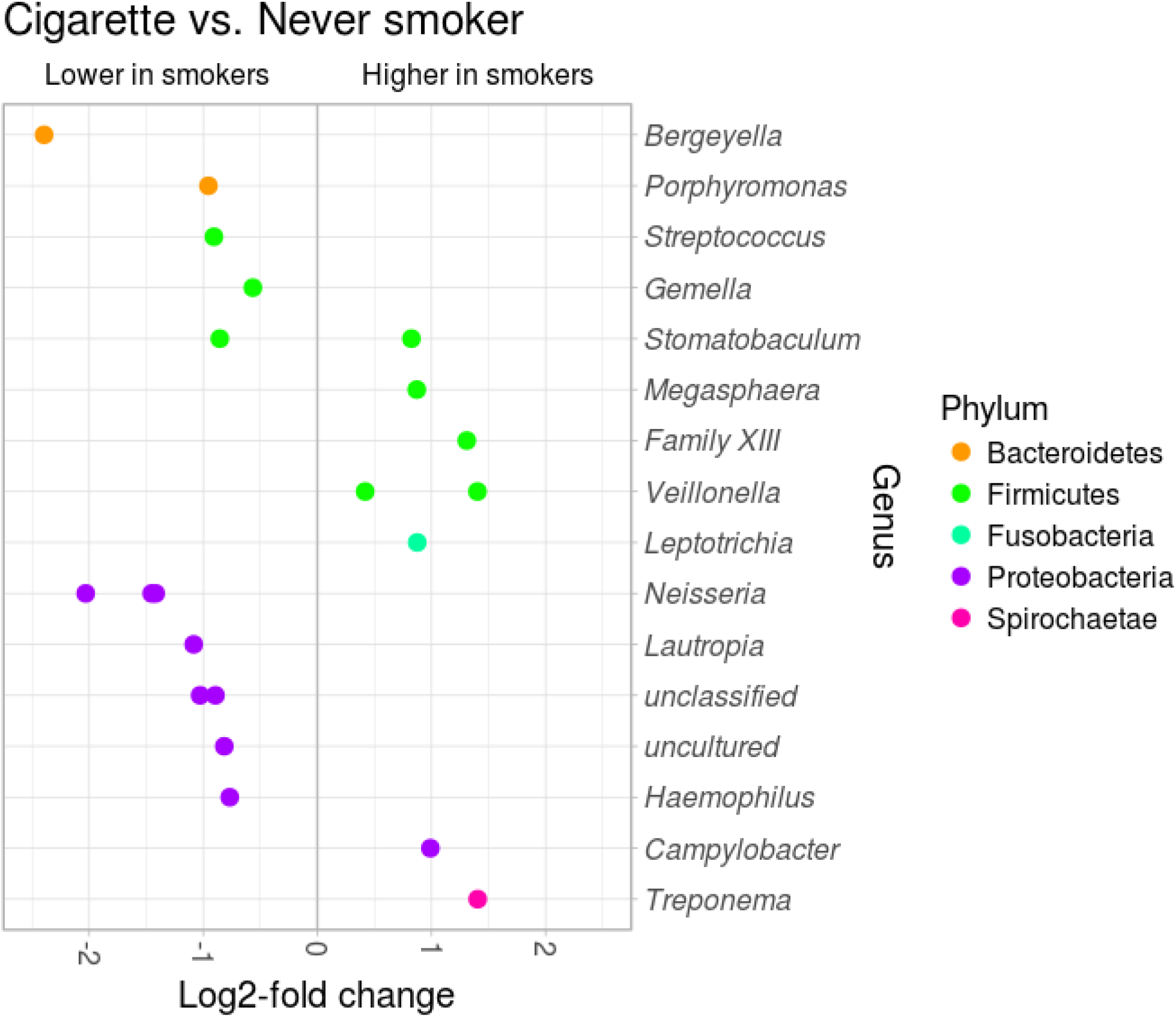
Adjusted multivariate differential analysis between current cigarette smokers (n=86) and never smokers (n=43). Starting from the 46 OTUs identified as differentially abundant from the crude model, adjusting for confounders OTU differentially abundant were reduced to 21.

**Figure 3:**
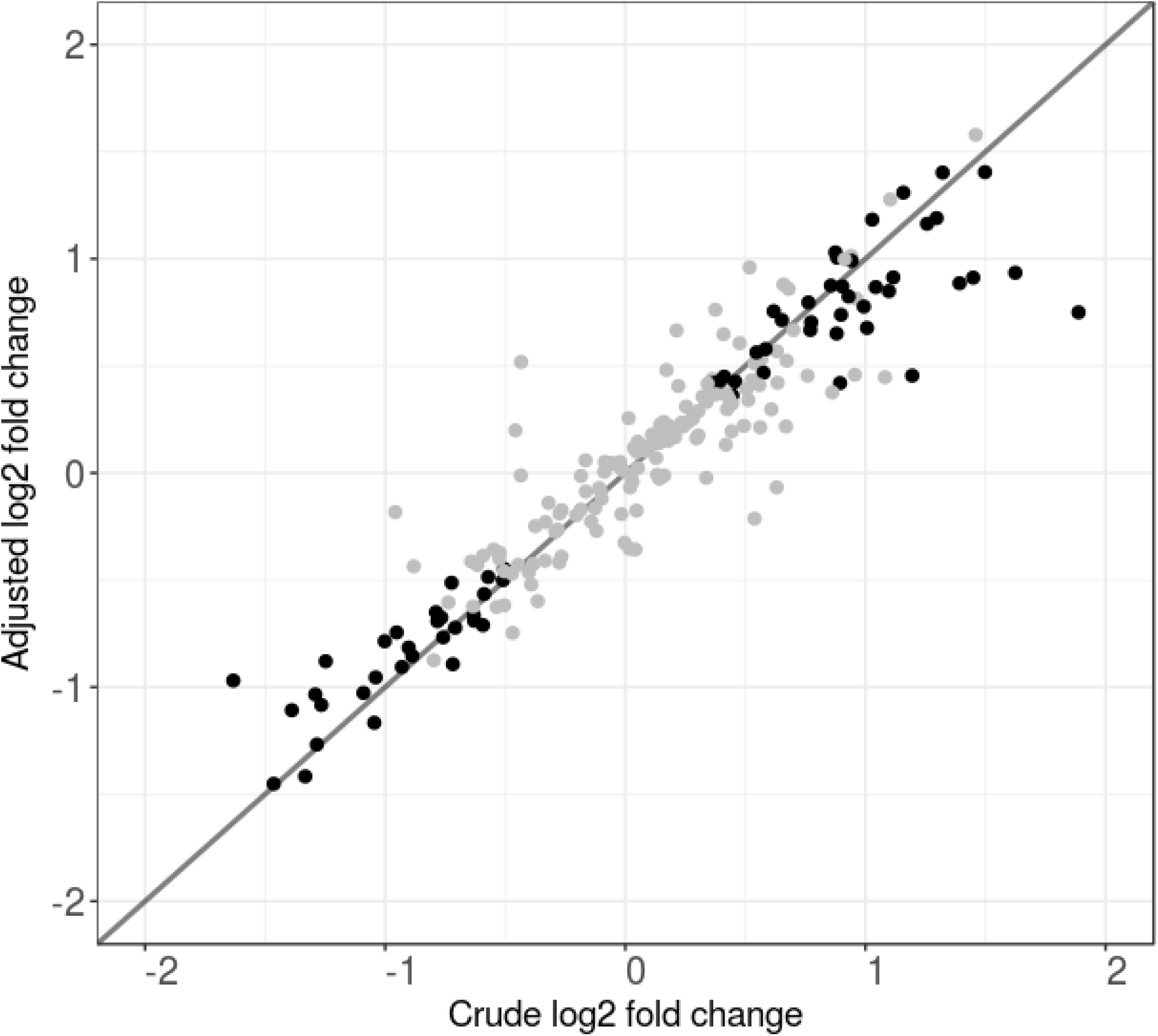
Comparison between 212 coefficients for current (n=86) vs. never smokers (n=43) from crude and adjusted negative binomial log-linear regression (adjusted for age, sex, race/ethnicity, self-reported physical activity, education, diabetes status, self-reported gum disease). Points in the scatter plot represent all differentially abundant OTUs, regardless of statistical significance, with black dot OTUs significant with the Wald test in crude analyses; coordinates are determined by the log2 fold change resulting from the crude analysis between current and never smokers (x axis) and the log2 fold change from the adjusted analysis between current and never smokers (y axis).

### Differences in oxygen utilization in the oral microbiome of smokers compared to never smokers

We functionally annotated the entire set of picked OTUs according to their oxygen requirement: 78 aerobic OTUs, 673 anaerobic OTUs and 395 facultative anaerobic OTUs. We failed to annotate 145 OTUs because their genera was annotated as uncultured bacteria or taxonomic resolution was higher than genus.

A minority of aerobic OTUs (29%) and a majority of anaerobic OTUs (60%) had higher mean abundance in current smokers as compared to never smokers. Facultative anaerobic OTUs were approximately evenly divided, with 51% having higher abundance in current smokers. We accordingly found differentially abundant OTUs between current smokers and never smokers to be over-represented in aerobic OTUs (Hypergeometric test, p = 0.004). Using Gene Set Enrichment Analysis (GSEA) to account for collinearity between OTUs and the direction of the abundance change (up / down), aerobic OTUs were significantly depleted among current smokers relative to never smokers (Enrichment Score test statistic ES = −0.75, p = 0.002, GSEA permutation test). Anaerobic OTUs were enriched in smokers relative to never smokers but the difference was not statistically significant (ES = 0.36, p = 0.14, GSEA permutation test). We also observed an enrichment of facultative anaerobic OTUs among never smokers compared to current smokers but this result was not statistically significant (ES = −0.29, p = 0.48, GSEA permutation test).

### Comparison of those with secondhand smoke exposure to never smokers

To provide independent evidence for causal inference, we compared the coefficients estimated for the contrast of current smokers versus never smokers to the coefficients for serum cotinine level, estimated from a non-overlapping group of self-reported nonsmokers (n=38) exposed to secondhand smoke. Consistency was assessed by calculating the Pearson Correlation of the two vectors of coefficients. This correlation is comparable to the Integrative Correlation Coefficient, which was originally proposed to assess the replicability of measurements from independent gene expression studies [60,61]. This correlation was estimated from the intersection of 121 OTUs passing the edgeR filter against low-variance features for both groups. Full differential abundance results for these 121 OTUs are reported in Supplemental File 1. We observed a positive correlation (Figure 4, Pearson’s Correlation = 0.40, p = 5e-6); this correlation was slightly attenuated after adjusted for age and educational attainment (Supplementary Figure 6, Pearson’s Correlation=0.28, p=0.002). This positive correlation identifies a similarity in the patterns of differential abundance in smokers vs. never smokers when compared to the shifts associated with increasing exposure to secondhand smoke. However, the application of three concurrent approaches for microbe set enrichment analysis (GSEA, GSVA, and ORA) on samples from participants exposed to secondhand smoke, with continuous or dichotomous serum cotinine levels as the response variable, did not identify significant enrichment or depletion of aerobiosis or anaerobiosis, reflecting smaller shifts associated with secondhand smoke exposure.

**Figure 4:**
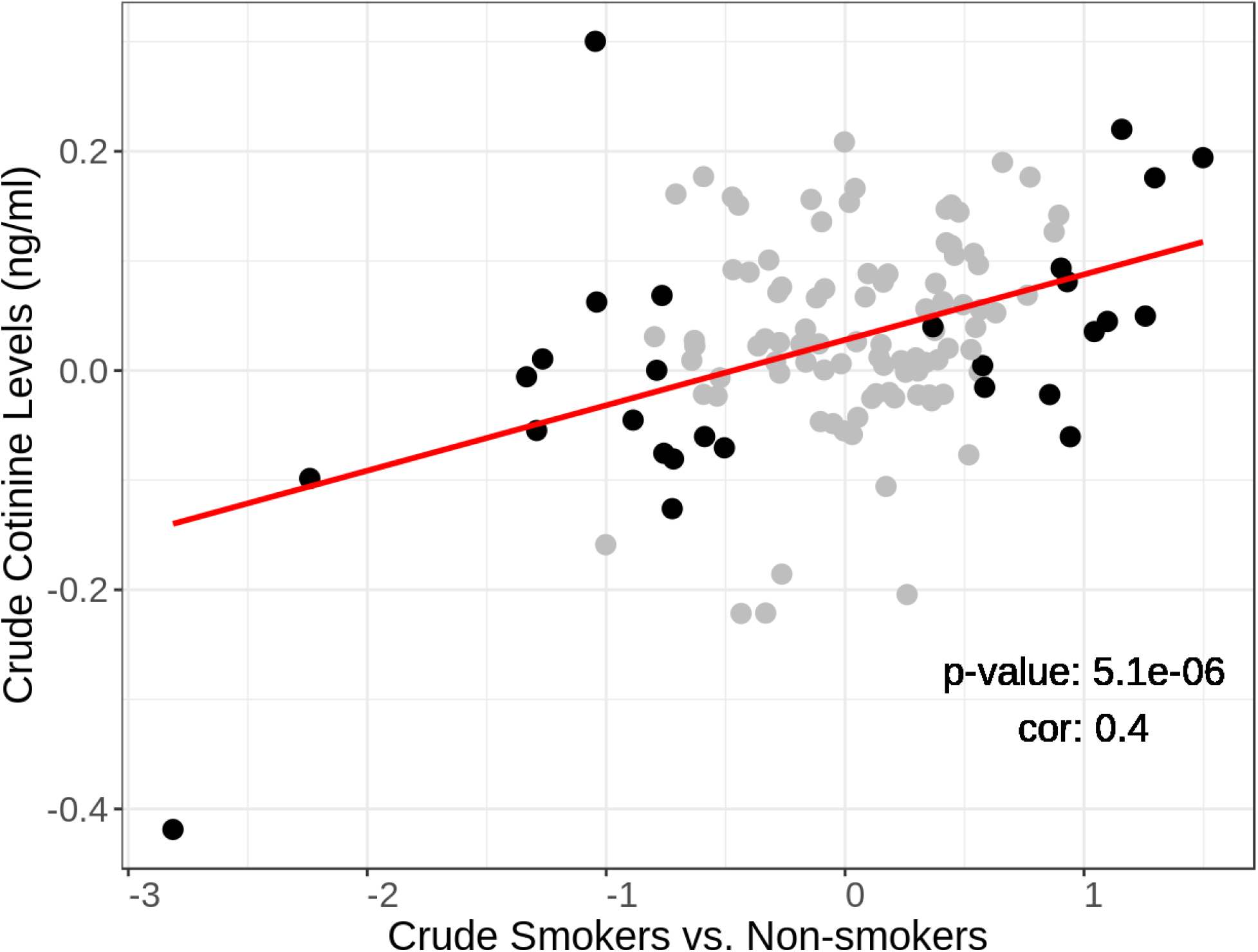
Comparison of log2 OTU fold changes between a) crude analyses of smokers (n=86) vs non-smokers with no detectable serum cotinine (n=43), and b) analysis of continuous cotinine levels among non-smokers exposed to secondhand smoke (n=38). Plot shows 121 OTUs that passed the edgeR filter for low-variance variables, including 28 OTUs that are differentially abundant in smokers (FDR <0.05, indicated in black). Coefficients for these two contrasts, involving different measures of exposure on different individuals, are positively correlated (Pearson Correlation = 0.58, p = 0.0013 for 28 OTUs; Pearson Correlation = 0.40, p = 5e-6 for all 121 OTUs)

### Comparison of alternative tobacco exposures to never smokers

Differential abundance of OTUs from participants who used e-cigarettes, hookah, and/or cigar/cigarillo but not cigarettes (Table 1) were contrasted to the never smoker group. Phyla Actinobacteria, Firmicutes and Proteobacteria were more abundant in alternative smokers while Bacteroidetes and an uncultured bacterium from Saccharibacteria were more depleted. In those who only smoked hookah (n = 28), genera *Porphyromonas, Leptotrichia, Streptobacillus, Fusobacterium*, and an uncultured bacterium from Saccharibacteria were depleted. No OTUs were identified as differentially abundant among users of e-cigarette (n=11) or cigar/cigarillo (n=23) who did not use any other smoking products. GSEA identified a significant depletion of aerobic OTUs in cigar and cigarillo smokers (ES = −0.697, p = 0.04 GSEA permutation test) and depletion of facultative anaerobic OTUs in e-cigarette and hookah smokers compared to never smokers (e-cigarettes ES = −0.514 p = 0.03 GSEA permutation test; hookah ES = - 0.489 p = 0.04 GSEA permutation test).

## Discussion

This study analyzes oral mouthwash specimens from a sub-sample of NYC HANES 2013-14 to provide multiple lines of causal inference supporting the hypothesis that tobacco smoke exposure alters the saliva microbiome. We found current smokers to harbour a different microbial composition compared to never smokers and the other tobacco exposure groups in terms of beta diversity, individual OTUs, and oxygen requirements. The microbiome of former smokers was more similar to never smokers than to current smokers. Phyla Candidate division SR1, Bacteroides and Proteobacteria were depleted in smokers with genera *Bergeyella, Porphyromonas, Prevotella, Haemophilus, Neisseria, Lautropia* and *Actinobacillus*, consistent with previous studies [62–64]. The depletion of Proteobacteria may be especially important as this depletion has also been found among individuals with periodontal disease compared to healthy controls [65]. Further, Proteobacteria levels in the oral microbiome have been associated with insulin resistance and inflammation [66]. These shifts in genera largely remained with adjustment for hypothesized confounders.

The large microbiome shifts associated with current smokers compared to never smokers included significant depletion of oxygen-requiring bacteria, and corresponding (but not statistically significant) enrichment of anaerobic bacteria. This finding is consistent with a proposed mechanism [67] by which smoking alters the oxygenation of the oral cavity, depleting oxygen and favouring anaerobic bacteria. Furthermore, the shifts in mean abundance occurring between current smokers and never smokers were positively correlated to those observed among non-smokers with varying levels of secondhand smoke exposure as measured by serum cotinine. This indicates a dose-response relationship for secondhand smoke exposure, and reduces the plausibility of residual confounding as an explanation for the shifts observed in these separate groups of participants.

Reduced aerobiosis and increased anaerobiosis in the oral cavity have implications for oral and systemic health. The Red Complex, a trio of anaerobic bacteria *(Treponema denticola, Porphyromonas gingivalis* and *Tannerella forsythia)* are linked with the development of periodontal disease [68]. While these primarily inhabit the dental plaque, an overall anaerobic oral environment may facilitate colonization. Although this study did not provide species-level resolution to observe the Red Complex, a previous study [69] found increased abundance of *Porphyromonas gingivalis* and *Tannerella forsythia* in the subgingival plaque of smokers compared to non-smokers. Furthermore, oral anaerobiosis could provide greater opportunity for movement of oral bacteria to distant anaerobic environments in the stomach and gut. This study demonstrates how oxygen utilization provides a simplifying measure that can be used by future studies of the oral microbiome and health.

In a mixed group of users of alternative smoking products including e-cigarettes, hookahs, cigars and cigarillos, we found some alterations comparable to those in cigarette smokers (like *Lactococcus* and *Neisseria* genera), while others like *Porphyromonas* had an opposite trend. This small and heterogeneous group of alternative smoking products does not allow robust conclusions, but indicates the possibility that alternative products could alter the oral microbiome composition in ways similar to cigarette smoke. As e-cigarettes are gaining popularity in use, [70] more research is needed to explore the effect of vaping on the oral microbiome.

This study has a number of limitations. As a cross-sectional study, changes to the oral microbiome in direct response to tobacco exposure were not measured; longitudinal data are needed to directly observe tobacco-induced changes to the oral microbiome. We defined secondhand exposure to smoke using a serum cotinine upper cut-off of 14 ng/mL [71], Four of 38 participants in the secondhand exposure group had serum cotinine higher than the cut-off value of 10 ng/mL [35], and may have misreported recent light cigarette usage. However, removing these four individuals had negligible effect on reported results. We adjusted for self-reported gum disease as a measure of periodontal health; this measure is imperfect and residual confounding by periodontal health may remain. Additionally periodontal health may be a mediator rather than a confounder. Regardless, self-reported gum disease was not strongly associated with beta diversity in our analyses, so adjusting for it should not impact results. Additionally, we did not adjust for differences in dietary habits given the general lack of validity of self-reported diet data [72]; we cannot rule out residual confounding by diet. However, our findings, which show a shift toward anaerobic bacteria among smokers make biological sense in response to smoke exposure, reducing the likelihood that these findings are the result of residual confounding. Finally, the current analysis is based on 16S rRNA gene analyses capturing only genus and higher-level taxonomic information; whole metagenomic sequencing may provide additional important information on shifts to the oral cavity caused by tobacco exposures and functional information.

The strengths of this study are that it included a racially/ethnically diverse group of participants and an array of tobacco exposure groups including self-reported non-smokers exposed to secondhand smoke. The study design allowed multiple, complementary comparisons, as well as biological analysis, to help distinguish causal associations from associations likely to be caused by residual confounding. This study introduces several analyses that are, to the best of our knowledge, novel to epidemiological studies of the human microbiome. These include: 1) the application of Gene Set Enrichment Analysis methods for biological interpretation of gross microbiome shifts, 2) use of a scatter plot to visualize the comparison of crude vs. adjusted regression coefficients in high-dimensional data, and 3) application of the Integrative Correlation Coefficient [60,61], a method introduced to assess reproducibility of gene expression studies, to make causal inference by comparing regression coefficients from different samples with different measures of tobacco exposure (smokers vs. non-smokers, and dose-response for continuous serum cotinine measurements).

## Conclusions

Overall shifts between aerobic and anaerobic microbiota is a relevant simplifying measure that should be considered in future health studies of the oral microbiome. These results support a plausible biological mechanism for population-level shifts in the oral microbiome caused by exposure to tobacco smoke, through three lines of observational evidence: 1) consistency of the microbiome shifts with reduced microbiota oxygen utilization as a biological mechanism for the shifts observed in smokers; 2) consistency of oral microbiome abundance fold-changes in current smokers versus non-smokers with abundance changes along the gradient of secondhand smoke exposure among non-smokers; and 3) tobacco-related associations that are stronger than associations with sociodemographic and health indicators, and that are not meaningfully affected by controlling for hypothesized confounders.

## Acknowledgements

We would like to thank Sharon Perlman and Jennifer Rakeman-Cagno from the New York City Department of Health and Mental Hygiene for their collaboration during study implementation and design.

## Appendix: Supplementary figures

**Supplementary Figure 1:**
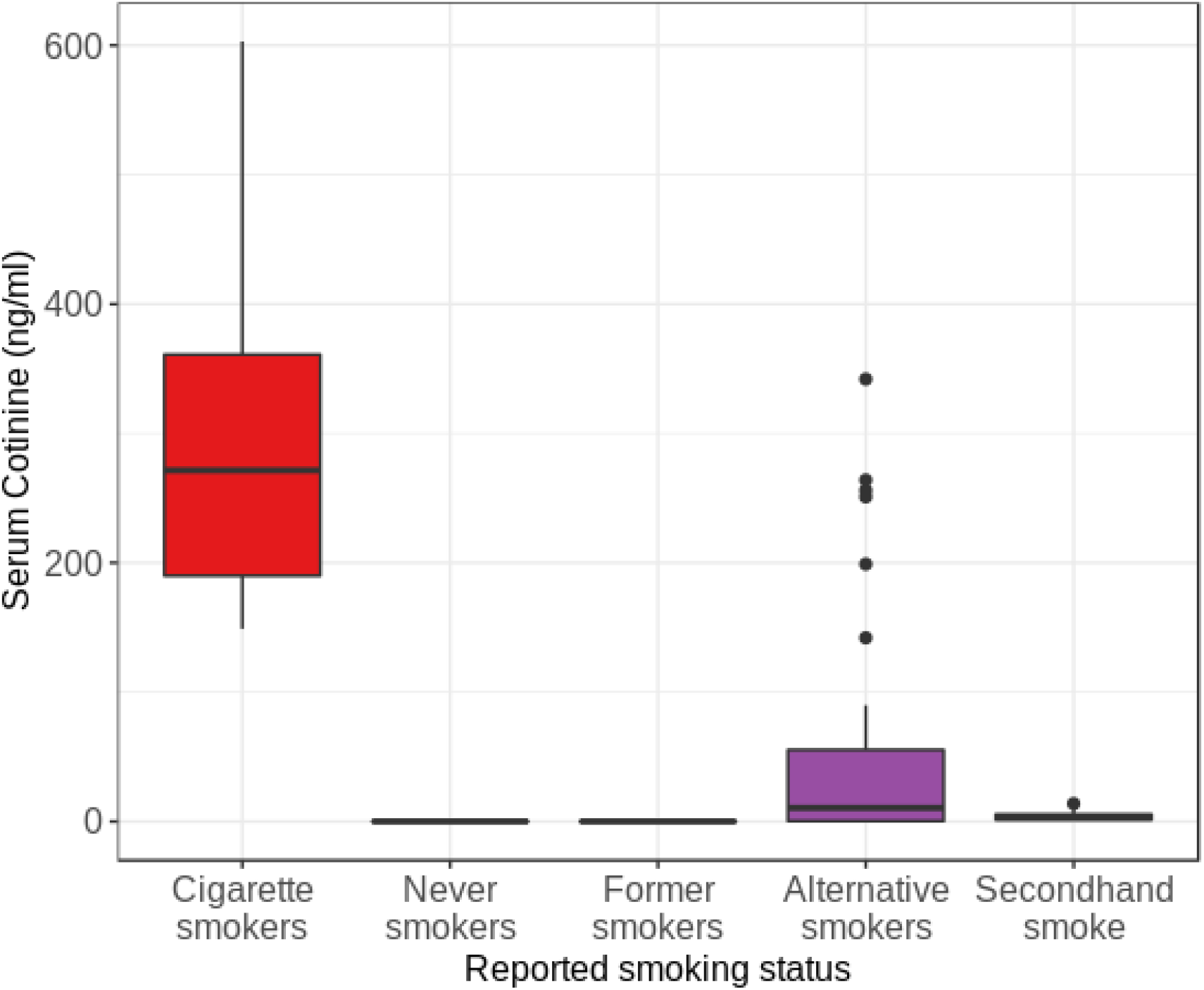
Distribution of measured serum cotinine among the five smoke exposure groups with current cigarette smokers followed by smokers of alternative tobacco products showing the highest serum cotinine levels.

**Supplementary Figure 2:**
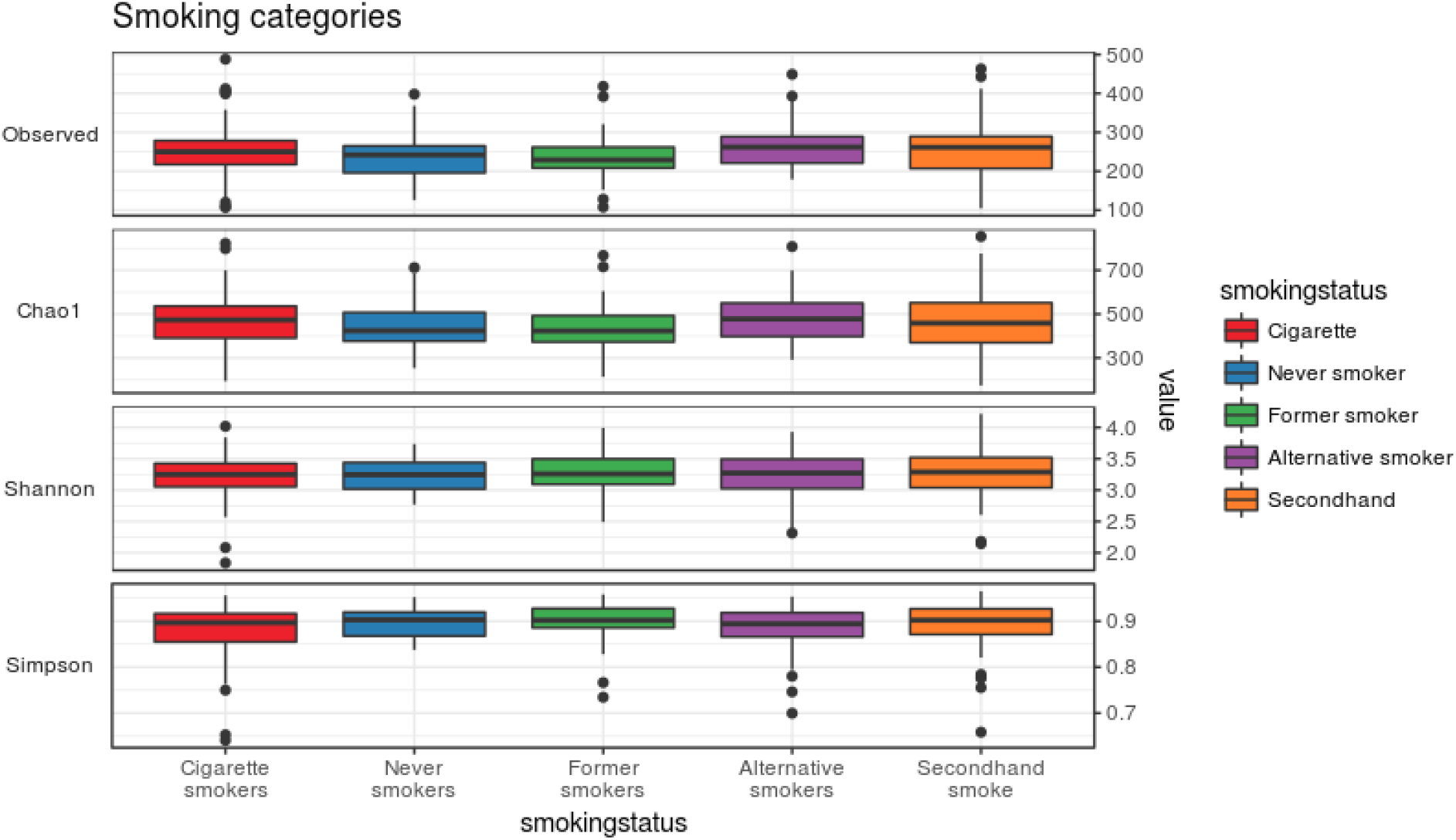
Alpha diversity measures between the five tobacco exposure groups. No significant differences were observed between groups with four measures of richness and evenness.

**Supplementary Figure 3:**
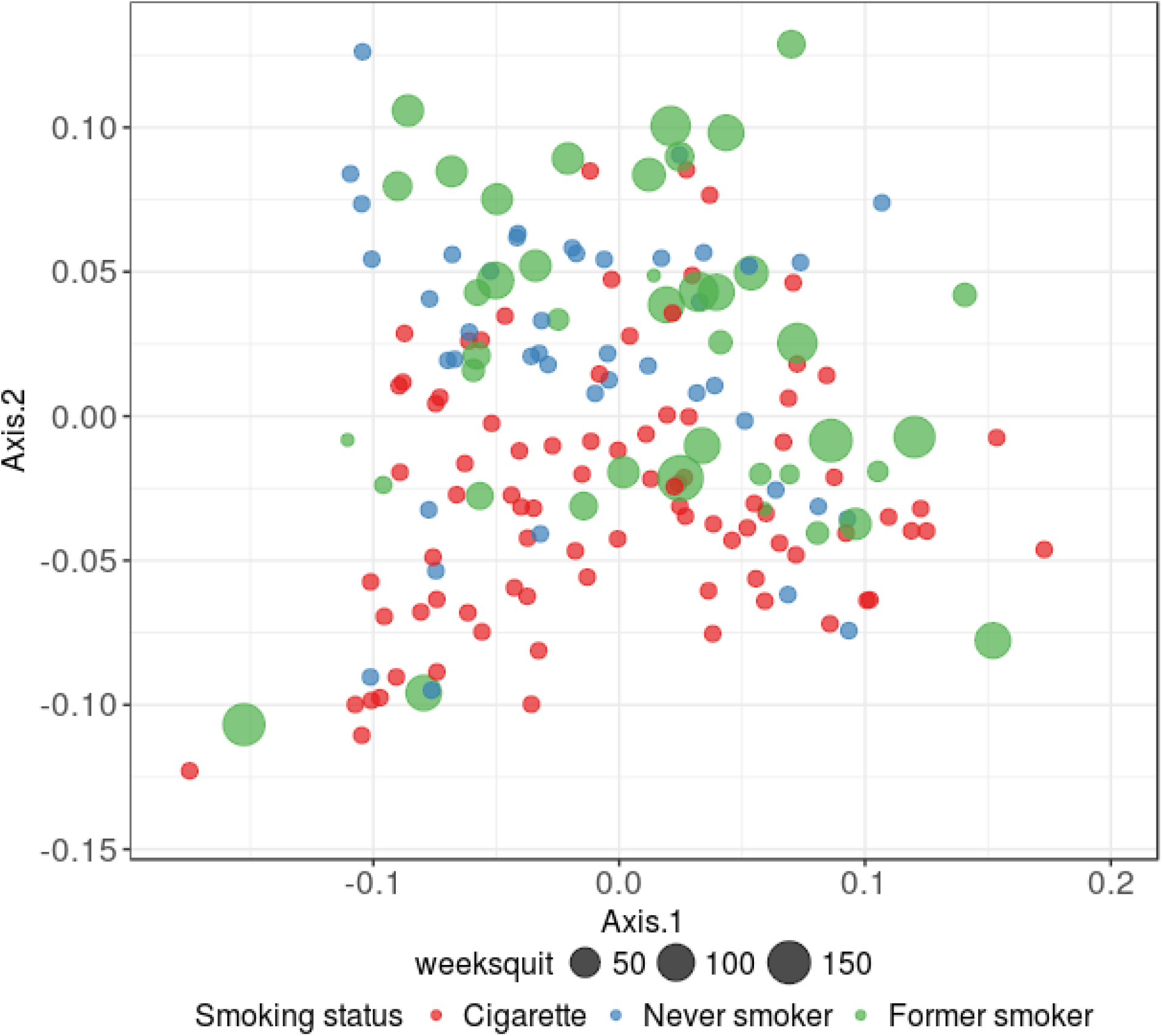
Principal coordinates analysis based on the weighted UniFrac distance on samples from cigarette smokers (n=86), never smokers (n=43), and former smokers (n=43) with size of dot indicating how long ago they reported quitting. A separation on the second axis between cigarette smokers and never smokers is present. We found former smokers were significantly different from current smokers (p=0.002, PERMANOVA test), but not from never smokers (p=0.16, PERMANOVA test).

**Supplementary Figure 4:**
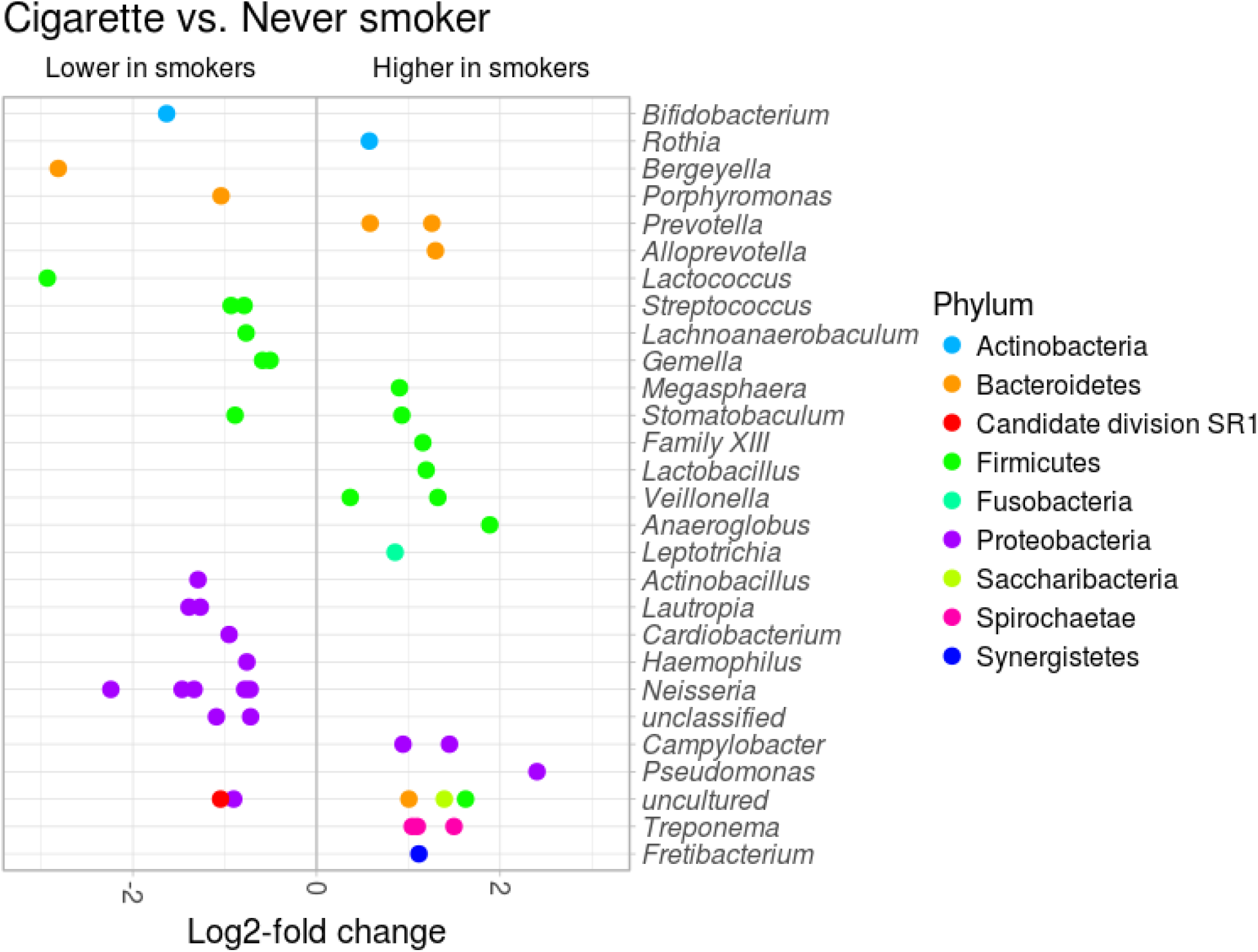
Crude differential analysis between current cigarette smokers (n=86) and never smokers (n=43). Dots in the plot are the 46 OTUs identified as differentially abundant without adjusting for hypothesized confounders and coloured according the taxonomy annotation at the phylum level. Position of the side of the plot is determined by the log2 fold change of abundance of the OTU.

**Supplementary Figure 5:**
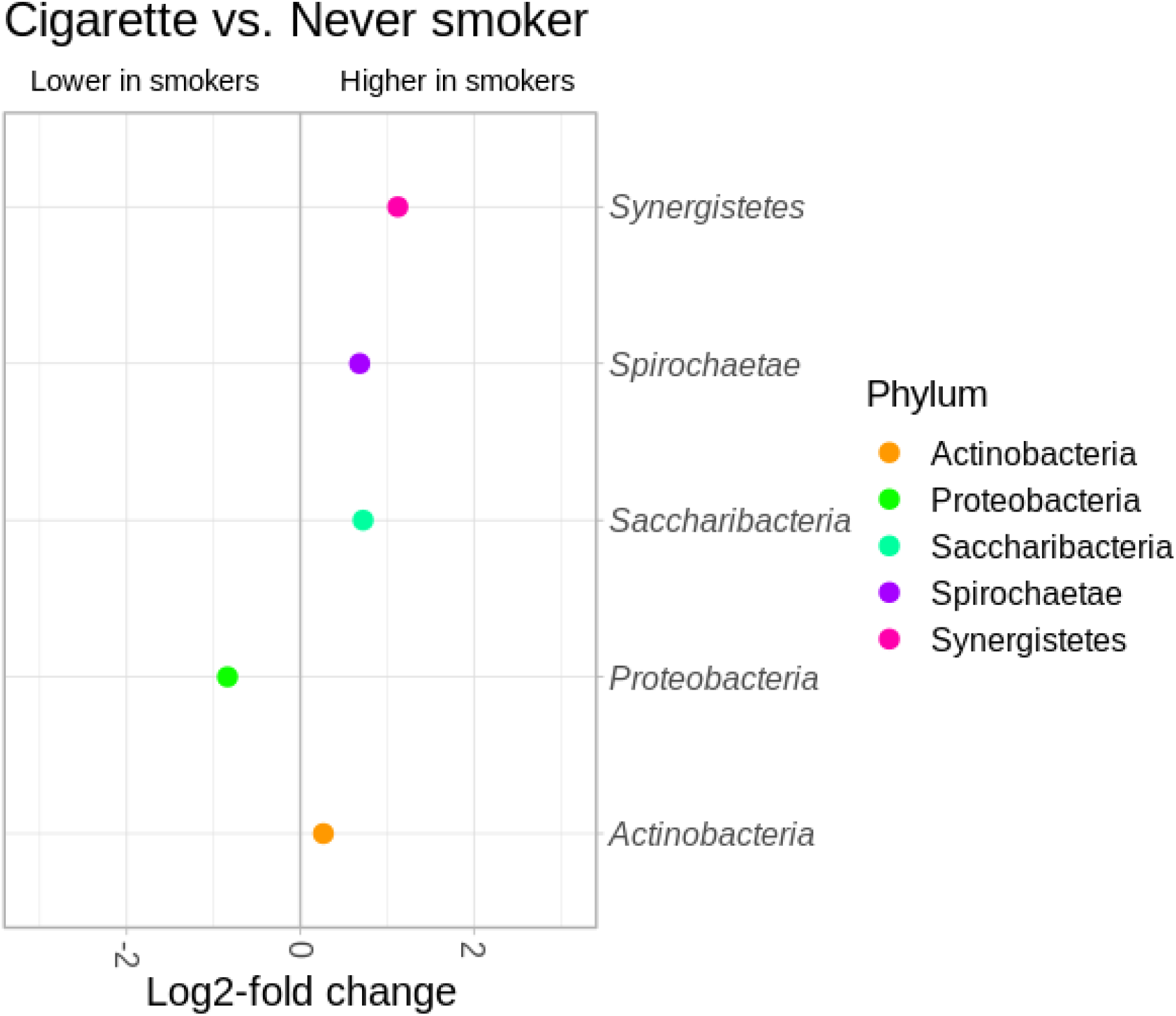
Crude differential analysis between current cigarette smokers (n=86) and never smokers (n=43) performed at the phylum level.

**Supplementary Figure 6:**
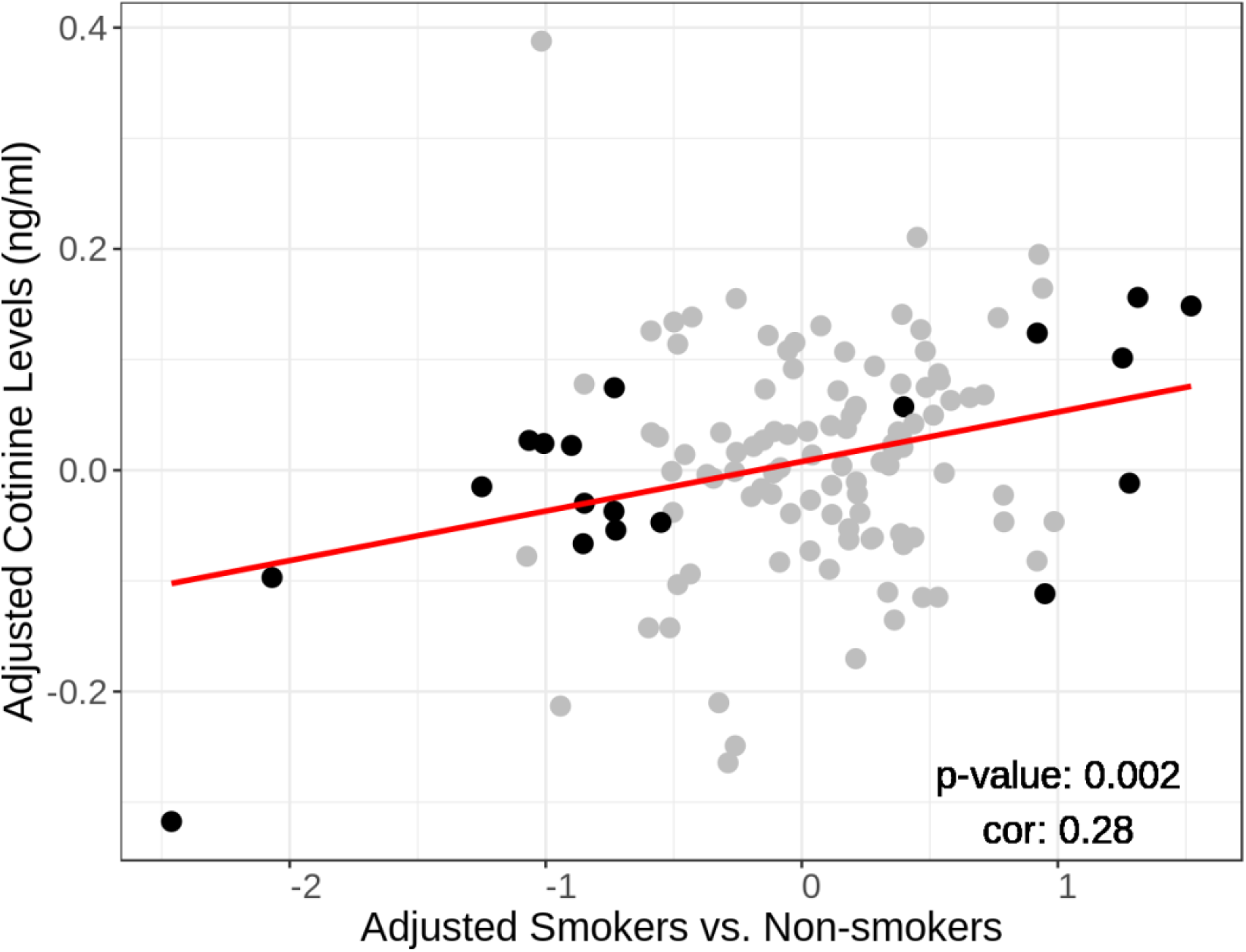
Equivalent of Figure 4, with adjustment for age and educational attainment. Comparison of log2 OTU fold changes between a) adjusted analyses of (x-axis) smokers (n=86) vs non-smokers with no detectable serum cotinine (n=43) and b) adjusted analysis of continuous cotinine levels among non-smokers exposed to secondhand smoke (n=38). Plot shows 121 OTUs that passed the edgeR filter for low-variance variables, including 19 OTUs differentially abundant in smokers (FDR<0.05, indicated in black). Similar to the crude model in Figure 4, a positive correlation was observed (Pearson Correlation = 0.68, p = 0.001 for 19 OTUs; Pearson Correlation = 0.28, p = 0.002 for all 121 OTUs).

## List of abbreviations

CI: - Confidence Interval
CUNY: - City University of New York
DNA: - Deoxyribonucleic Acid
ES: - Enrichment Score
FDR: - False Discovery Rate
GSEA: - Gene Set Enrichment Analysis
GSVA: - Gene Set Variation Analysis
HMP: - Human MIcrobiome Project
IRB: - Institutional Review Board
NHANES: - National Health Nutrition and Examination Survey
NYC DOHMH: - New York City Department of Health and Mental Hygiene
NYC HANES: - New York City Health Nutrition and Examination Survey
OR: - Odds Ratio
ORA: - Over-representation Analysis
OTU: - Operational Taxonomic Unit
PCOA: - Principal Coordinates Analysis
PERMANOVA: - Permutational Multivariate Analysis of Variance
RNA: - Ribonucleic Acid
rRNA: - Ribosomal Ribonucleic Acid

